# Tie-lines reveal interactions driving heteromolecular condensate formation

**DOI:** 10.1101/2022.02.22.481401

**Authors:** Daoyuan Qian, Timothy J. Welsh, Nadia A. Erkamp, Seema Qamar, Jonathon Nixon-Abell, Georg Krainer, Peter St George-Hyslop, Thomas C. T. Michaels, Tuomas P. J. Knowles

## Abstract

Phase separation of biomolecules give rise to membraneless organelles that contribute to the spatiotemporal organisation of the cell. In most cases, such biomolecular condensates contain multiple components, but the manner in which interactions between components control the stability of condensates remained challenging to elucidate. Here, we develop an approach to determine tie-line gradients in ternary liquid-liquid phase separation (LLPS) systems, based on measurements of the dilute phase concentration of only one component. We show that the sign of the tie-line gradient is related to the cross-interaction energy between the polymers in the system and discriminates between competitive and cooperative phase separation. Using this approach, we studied the interaction between protein Fused in Sarcoma (FUS) and polyethylene glycol (PEG) polymer chains, and measured positive tie-line gradients. Our results show that PEG drives LLPS through an associative interaction with FUS and is not an inert crowder. We further studied the interaction between PolyA RNA (3.0±0.5kDa) and the protein G3BP1, and using the tie-line gradient as a reporter for the stoichiometry of polymers in the condensate we determined a G3BP1-to-PolyA RNA molar ratio of 1:4 in the dense phase. Our framework for measuring tie-line gradients opens up a route for the characterisation of interaction types and compositions in ternary LLPS systems.

## I. INTRODUCTION

Biomolecular condensates play important roles in cells, both in healthy physiological function and disease development [1–5]. These condensates form via liquid-liquid phase separation (LLPS), driven by interactions between biomolecules such as protein and RNA [6, 7]. LLPS is often studied by generating phase diagrams that show under what conditions condensates form. However, we found phase diagrams alone contain insufficient information on whether solutes co-localise in the condensate or prefer to be in separate phases, and the concept of tie-lines becomes important. A tie-line is defined in the following manner. Take a ternary LLPS system with two solutes - a protein and an agent - and induce phase separation by mixing a suitable amount of each. The dilute and dense phases contain certain concentrations of the protein and the agent, and we can plot the two phases as two points on the phase diagram with concentrations as their coordinates. The tie-line is the line connecting these two points. This line indicates if the solutes prefer to be in the same or different phases, which gives a positive or negative tie-line gradient respectively. Using the Flory-Huggins model we found that tie-line directions directly relate to the cross-interaction between solutes. In the case of a positive gradient we can further approximate the gradient as the stoichiometry of solutes in the dense phase. Tie-line measurements have been performed on complex coacervates [8] by directly measuring concentrations of both solutes in dilute and dense phases, however such a measurement is challenging for protein condensates.

In this paper we demonstrate tie-line gradients can be determined by measuring the concentration of only one solute in the dilute phase. We do not require measurement of the other solute or dense phase concentrations, both enabling investigations with solutes that cannot be easily labelled or measured and ensuring minimal artificial effect from attaching florescent tags to LLPS-inducing agents [9]. Here we studied three systems. The first and second involve the protein Fused in Sarcoma (FUS). FUS is a 75kDa nucleic acid binding protein known to be key for controlling gene expression across cell types [10, 11] and closely related to the neurodegenerative disease amyotrophic lateral sclerosis (ALS) [12]. FUS is also known to undergo LLPS in response to many stimuli including salt [13], RNA [14], and polyethylene glycol (PEG) – where PEG was previously assumed to be an inert crowder [12, 15]. We investigated LLPS of FUS with either PEG20k (20kDa) or PEG10k (10kDa). In both systems we found FUS and PEG co-localise in condensates, meaning PEG does not act as a crowder. Our third system involves the protein G3BP1, a 60kDa GTP-ase activating protein which has RNA-binding properties. G3BP1 is known the be the core protein responsible for nucleating the formation of stress granules [6, 16], a type of membrane-less organelle responsible for controlling stress response in cells [1, 17], and G3BP1 is also associated with cancer development [18, 19]. As part of its role in stress granule formation, G3BP1 forms favourable interactions with RNA to induce formation of condensates [1]. We studied the interaction between G3BP1 and single stranded PolyA RNA (3.0±0.5 kDa) and found that G3BP1 and the PolyA colocalize in the condensate with a molar ratio of 1:4. Our method can be widely applied to characterise interactions and compositions of condensates in other LLPS systems too.

## II. SOLUTE INTERACTION DIRECTLY RELATES TO TIE-LINE GRADIENT

Here we use the Flory-Huggins model to investigate tie-lines. The Flory-Huggins free energy density *f* of mixing a solvent with volume fraction *ϕ*_0_, and two different polymer solutes with lengths *N*_1_, *N*_2_ and volume fractions *ϕ*_1_, *ϕ*_2_ respectively, is [20–23]

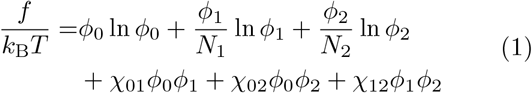

with *k*_B_*T* the unit thermal energy, *χ*_*µν*_ the effective interaction between components *µ* and *ν*, and the solvent volume fraction is constrained by the condition *ϕ*_0_ = 1 − *ϕ*_1_ − *ϕ*_2_. The first three terms in equation (1) are entropic contributions that prefer mixing, while the remaining terms are interactions that can favour either mixing or phase separation, depending on values of *χ*_*µν*_. Taken together, the free energy density landscape *f* (*ϕ*_1_, *ϕ*_2_) can be used to deduce phase-separation propensity at a particular combination of solute concentrations (*ϕ*_1_, *ϕ*_2_): if the free energy density is concave in any direction, the system is susceptible to thermal fluctuations and will spontaneously phase separate, this region of the phase diagram is the spinodal; on the other hand, a convex free energy density traps the system in a ‘valley’ and the mixture is locally stable. Outside the spinodal region, however, phase separation is still possible if multiple free energy minima exist and this region of metastability is denoted the binodal. A definition of tie-line gradient is natural once LLPS occurs. Denote dilute and dense phase concentrations of the two solutes as 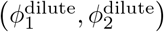 and 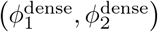, the gradient of the tie-line is simply 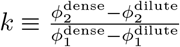 since it connects both points. In the case where both solutes have low dilute phase concentrations we approximately have 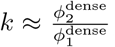, i.e. the volume ratio of solutes in the dense phase.

We use a minimal model (Appendix A 1) to illustrate the phase space structure of the Flory-Huggins system. Setting solutes to unit length *N*_1_ = *N*_2_ = 1 and assuming inert solvent *χ*_01_ = *χ*_02_ = 0, the only relevant parameter is *χ*_12_ ≡ *χ*. We calculate the spinodal region analytically using the Hessian and compute the binodal and tie-lines numerically via convexification of *f* (*ϕ*_1_, *ϕ*_2_) [24]. These two boundaries intersect and are parallel to one another at critical points. The complete phase space of the minimal model is characterised by a cooperative branch (where both solutes are enriched in one phase) at *χ <* −8 with positive tie-lines and a competitive branch (where solutes are enriched in separate phases) at *χ >* 2 with negative tie-lines (figure 1). It is important to note that the phase boundaries for competitive and cooperative phase separation in the dilute regime bear great semblance to each other, so measurements of phase diagram shapes in small portions of phase space cannot determine whether cooperative or competitive interactions are drivers of LLPS.

**FIG. 1.**
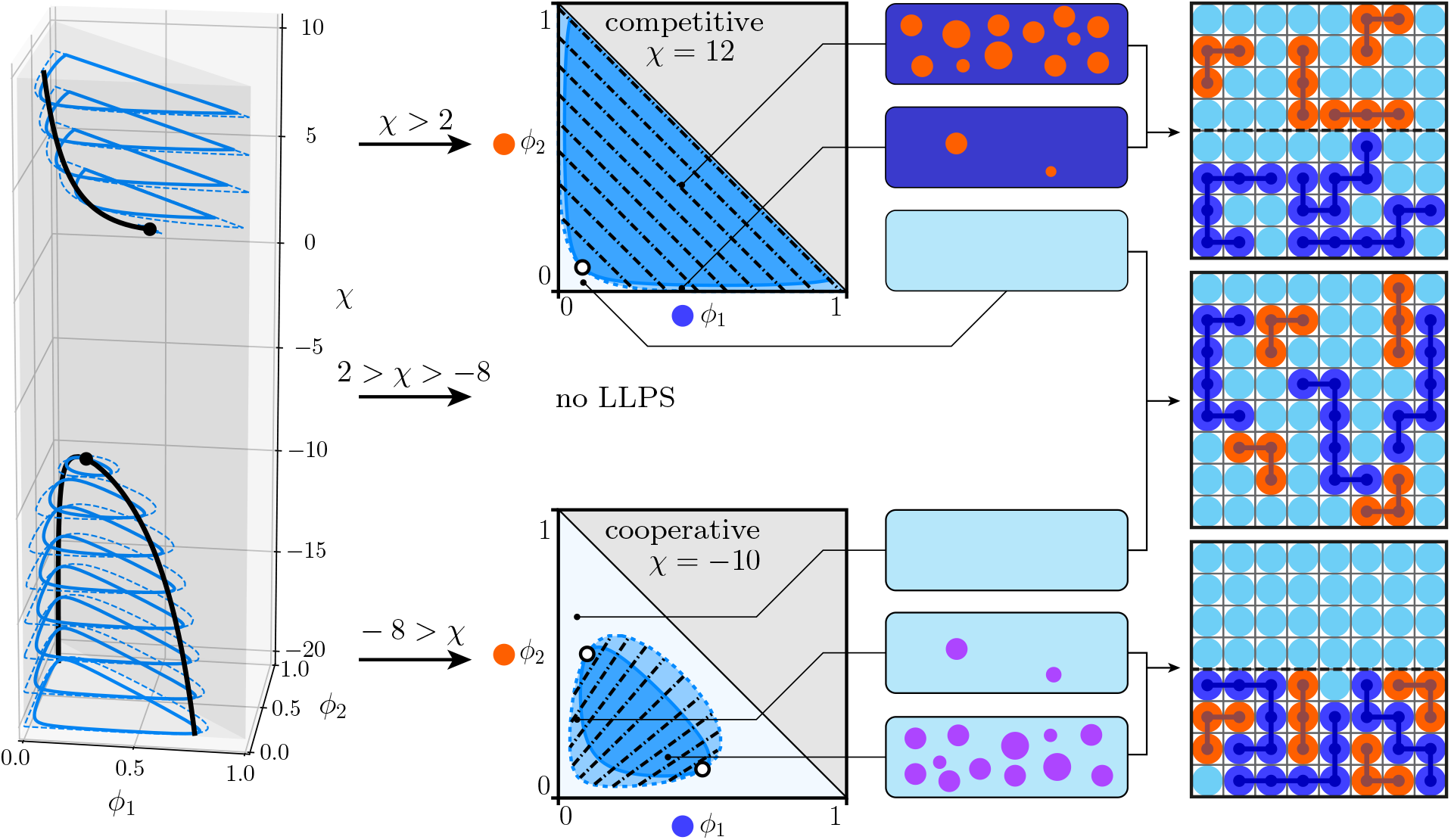
Phase space structure of a ternary system. Left column: critical points (black dots and black lines), spinodal boundaries (blue solid lines) and binodal boundaries (blue dashed lines) in the (*ϕ*_1_, *ϕ*_2_, *χ*) space. No phase separation occurs in the 2 *> χ >* −8 range. Central column: cross-sections of phase space at *χ* = 12 (top) and *χ* = −10 (bottom), corresponding to phase diagrams with critical points (hollow circles), spinodal (dark blue region), binodal (light blue region) and tie-lines (black dashed lines) plotted. In the competitive case the individual solutes de-mix into separate compartments while in the cooperative case they co-localise in condensates. Right column: lattice model illustration of the LLPS configurations.

To derive an approximate expression for the tie-line gradient *k* we use the full Flory-Huggins expression and diagonalise the Hessian. We approximate *k* as the gradient of the eigenvector with the smaller eigenvalue, which corresponds to the locally preferred direction of phase-separation (Appendix A 2). We then show that the sign of *k*, sgn(*k*), is sgn(*k*) = −sgn (1 + 2*χ*^Δ^) with 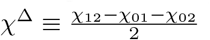. Assuming a dilute *ϕ*_1_ ≪ 1 we further obtain

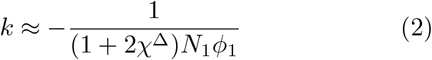

To visualise the physical interpretation of *χ*^Δ^ we recall the definition of the original 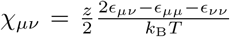 with *ϵ*_*µν*_ the bare contact energy between *µ* and *ν* particles and *z* a coordination constant. Direct substitution gives 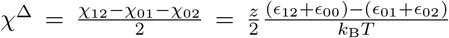. It is evident that *χ*^Δ^ signifies the energy difference between forming cooperative and competitive phases. We compare binodal tie-lines and local Hessian eigenvectors in the dilute regime of the phase diagram (figure 2) and note the two agree with each other relatively well.

**FIG. 2.**
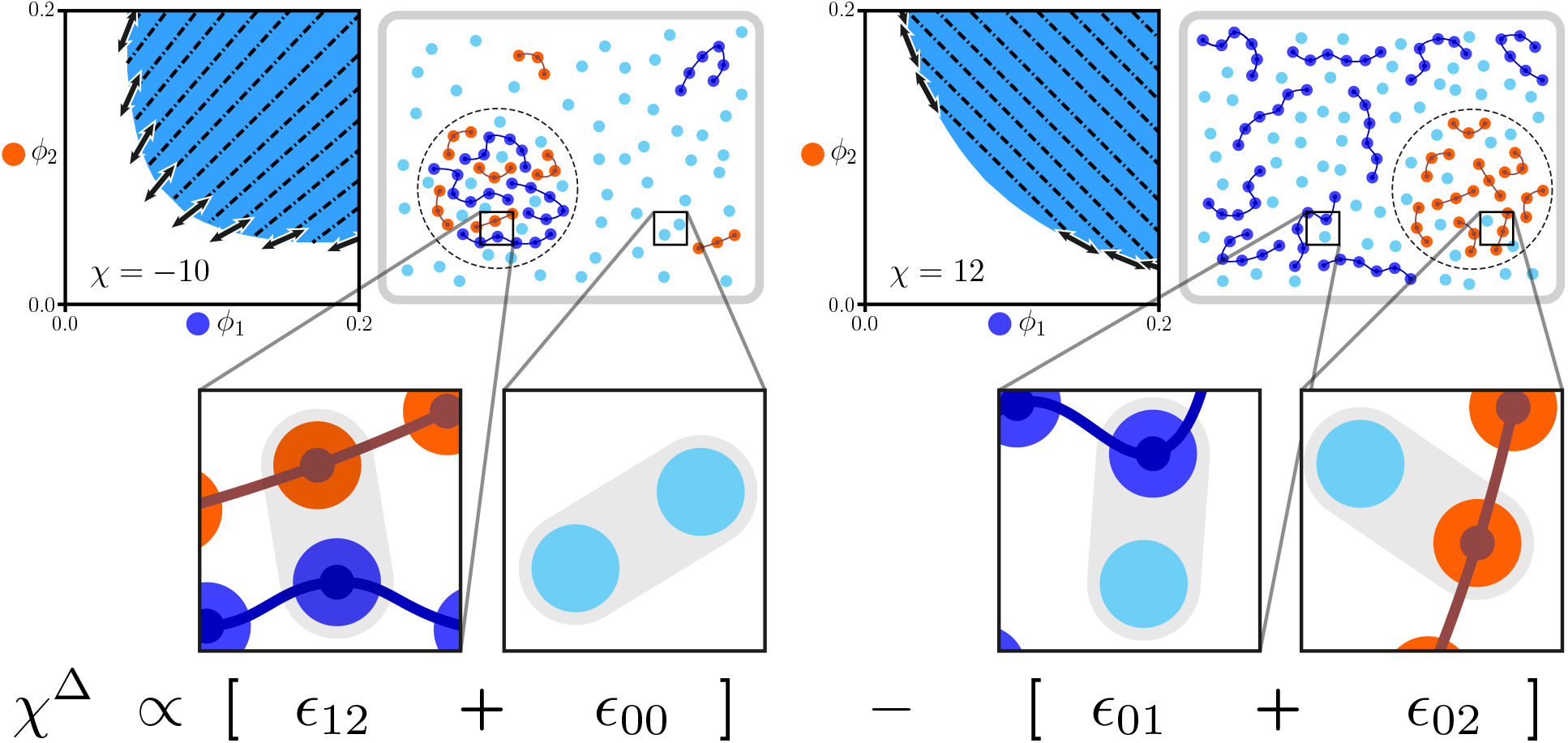
Sign of the tie-line gradient is determined by the cross interaction energy *χ*^Δ^. Top: comparison between tie-lines generated from numerical convexification (black dashed lines) and from diagonalising the Hessian (black thick arrows), with graphical illustrations of partitioning of solutes and solvent. Dark blue regions are the binodal. Bottom: graphical illustration of the physical interpretation of the *χ*^Δ^ parameter. *χ*^Δ^ is the energy difference between forming (left) cooperative phases and (right) competitive phases. Notice homotypic interactions *ϵ*_11_ and *ϵ*_22_ do not enter *χ*^Δ^, although they still affect the magnitude of the tie-line gradient.

## III. Results AND DISCUSSION

### Measurement approach

Measuring tie-lines using the dilute phase concentration of one component relies on the fact that a tie-line, by definition, is a line on which two points in the phase-separated space give the same dilute and dense phase compositions. As such, the problem reduces to finding two points in the phase diagram within the LLPS region that gives the same dilute phase concentration of one of the components - say *ϕ*_1_ - assuming the phase boundary has some finite gradient. To locate two such points we can first pick a point (the ‘anchor’) in the phase-separated region at 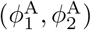 and measure the dilute phase concentration of *ϕ*_1_ as 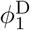. We then pick a series of points (the ‘linescan’), keeping the total *ϕ*_1_ concentration constant at some other value 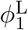 while varying the *ϕ*_2_ concentration. We can then interpolate the linescan measurements and find the point with the same dilute phase concentration 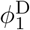. This pair of points allows us to construct the tie-line by connecting them with gradient *k* calculated as 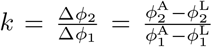 (figure 3A). Experimentally, we attached florescent tags to FUS and G3BP1 proteins and used a home-built confocal setup [25] to determine the photon count intensity from the dilute phase, which directly relate to the protein concentration (the conversion is not necessary since the 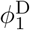 value does not enter the tie-line expression); a microfluidic device was used to trap the phase separating systems in water-in-oil droplets so that measurements could be carried out without surface effects (figure 3B, 3C and Appendix B 1).

**FIG. 3.**
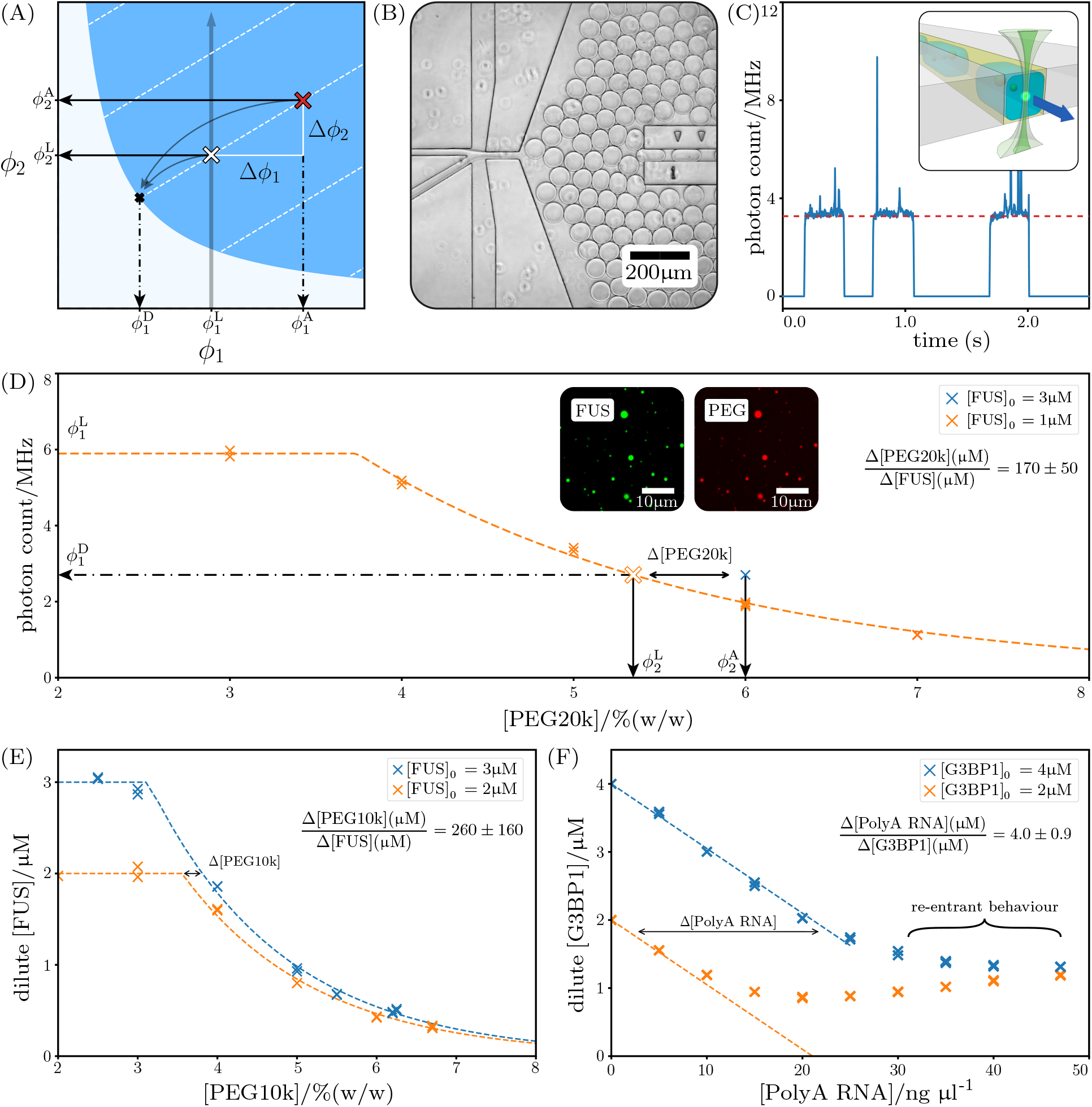
(A) Illustration of anchored linescan. Blue region is the binodal with representative tie-lines (white dashed lines). The anchor is plotted as a red cross and linescan plotted as a vertical grey solid line. By comparing the dilute phase *ϕ*_1_ concentrations 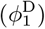 one identifies from linescan the second point on the tie-line (white cross). The gradient can then be calculated as 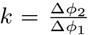. (B) Bright-field microscopy image of the microfluidic device. Droplets are generated on the left and pushed into a narrow channel on the right. Measurements were performed at the end of the channel to allow sufficient incubation time. (C) Raw signal coming from three droplets (blue solid line), characterised by plateaus from dilute phase readings and spikes from condensates passing through the confocal spot. During analysis a 3-*σ* cut-off was used to filter out the spikes to give an estimate of the dilute phase photon count, averaged over ∼100 droplets (red dashed line). Regions of low photon counts are gaps between droplets. Inset: 3D illustration of the channel cross-section and the confocal profile. (D) Results of anchored linescan for the FUS-PEG20k system. Solid crosses are measured values and the hollow cross is interpolated. The dashed line is a phenomenological fit. Measurement errors are comparable to marker sizes. Inset: Confocal microscopy images of condensates with both FUS and PEG20k labeled. (E) Serial linescans for the FUS-PEG10k system. Dashed lines are phenomenological fits. (F) Serial linescans for G3BP1-PolyA RNA system, showing an initial drop in dilute phase [G3BP1] due to phase separation and a rise at higher [PolyA RNA] due to the re-entrant effect. Dashed lines are linear low-[PolyA RNA] fits. Legends in (D) - (F) give total starting protein concentrations in droplets.

The anchored linescan is the minimal measurement needed to determine the tie-line gradient and we applied this to the FUS-PEG20k system. We then extended the method by performing multiple linescans and compared dilute phase concentrations between these, this is done in the FUS-PEG10k and G3BP1-PolyA RNA systems (Appendix B 2). Proteins are produced with an insect cell line (Appendix B 3) and a summary of molecular weights, densities and polymer lengths of individual components used in the following calculations are listed in Appendix B 4.

### Anchored linescan with FUS-PEG20k

We chose the anchor at 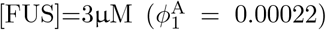 and 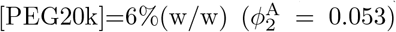. The linescan was performed at 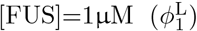 for [PEG20k] in the range of 3.0∼7.0%(w/w). This gives Δ[FUS]=2µM. The linescan data was then fit to a phenomenological curve and we extract Δ[PEG20k]= (0.65±0.20)%(w/w) (Appendix B 5). Thus we obtain the tie-line gradient 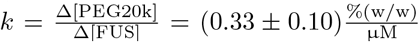 (figure 3D). Converting to molar ratio we get *k* = 170±50. Volume and mass ratios are calculated and summarised in table I.

**TABLE I.** Summary of ratios of components in condensates in different units and cross-interaction energies *χ*^Δ^. PEG is the main polymeric component in FUS-PEG condensates in molar, volume and mass ratios while in G3BP1-PolyA RNA condensates PolyA dominates in number, and G3BP1 dominates in volume and mass. *χ*^Δ^ estimates show FUS-PEG and G3BP1-PolyA interactions are weakly and strongly attractive respectively.

|  | FUS : PEG20k | FUS : PEG10k | G3BP1 : PolyA |
| --- | --- | --- | --- |
| Molar ratio | 1 : $170 \pm 50$ | 1 : $260 \pm 160$ | 1 : $4.0 \pm 0.9$ |
| Volume ratio | 1 : $38 \pm 11$ | 1 : $29 \pm 18$ | 1 : $0.12 \pm 0.03$ |
| Mass ratio | 1 : $32 \pm 9$ | 1 : $24 \pm 15$ | 1 : $0.14 \pm 0.04$ |
| $\chi^\Delta$ | -0.45 | -0.74 | -70* |

\* Flory-Huggins theory breaks down

The resulting positive tie-line is surprising, since we expected PEG to act as an inert crowder [12, 15] and it should thus have a higher concentration in the bulk to exert a positive net partial pressure on the condensates. We verified our results qualitatively by preparing condensates with fluorescently labeled FUS and PEG20k. Confocal microscopy (Appendix B 6) showed that condensates are both richer in FUS and PEG20k in comparison to the dilute phase, confirming our finding that FUS and PEG20k form condensates cooperatively. Furthermore, since 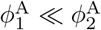 we use equation (2) to estimate *χ*^Δ^ ≈ − 0.45 so FUS and PEG20k are weakly attractive, with an effective interaction of half of *k*_B_*T* between lattice sites.

### Serial linescans with FUS-PEG10k

To confirm that the positive tie-line gradient is typical for PEG and not a coincidence due to the length of choice (20kDa), we measured the tie-line gradient for FUSPEG10k using serial linescans without anchor (figure 3E). Furthermore we also did a calibration series to map intensity to actual concentrations (appendix B 1). The tie-line gradient can be extracted (Appendix B 5) to give 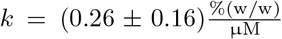 and molar ratio *k* = 260±160, on the same order of magnitude as the PEG20k result, indicating there is a similar cooperative interaction between FUS and PEG10k. Using equation (2) at [FUS]=1µM and [PEG10k]=5%(w/w) we obtain *χ*^Δ^ = −0.74. We further used the linescan data to reconstruct the binodal boundary and tie-lines, and fitted the resulting phase diagram to the Flory-Huggins theory by generating numerical phase diagrams using the convex hull algorithm. The fitting gives *χ*^Δ^ ≈ − 0.54 (Appendix C). Moreover, we tested dextran, another carbohydrate polymer, and also found a cooperative interaction with FUS (Appendix D).

### Serial linescans with G3BP1-PolyA RNA

We used single-stranded PolyA RNA (3.0±0.5 kDa) to form protein-RNA condensates with G3BP1 and carried out serial linescans (figure 3F). The low-[PolyA RNA] region shows a steady decrease in dilute phase [G3BP1] due to phase separation, and curves up at higher [PolyA RNA] as it enters the re-entrant branch [26, 27], where the additional PolyA RNA contributes to condensate dissolution instead of formation. We performed a linear fit for the low-[PolyA RNA] branch and the tie-line gradient is 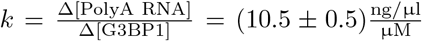, corresponding to a molar ratio of *k* = 4.0 ± 0.9. This indicates a clear associative interaction between the G3BP1 and PolyA RNA. Using equation (2) to extract *χ*^Δ^ however leads to a contact energy of the order -70*k*_B_*T* at concentrations [G3BP1]=4µM and [PolyA RNA]=20ng/µl. This is clearly unphysical and the large magnitude likely arises from underestimation of the dense phase interaction energies in the Flory-Huggins picture. G3BP1 is an RNA-binding protein and once bound, there is strong spatial correlation between the molecules and adopting a mean-field picture inevitably weakens the interactions so a much larger magnitude of *χ*^Δ^ is needed to compensate.

### Results summary

Table I summarises the molar, volume and mass ratios of the components in condensates as well as estimated cross-interaction energies. For the FUS-PEG systems the main polymeric component in condensates, by all 3 metrics, is PEG. This highlights a potential caveat in using PEG to induce LLPS in vitro: the high PEG content in condensates can affect the behaviour of proteins as compared to in vivo condensates, so functional conclusions derived from in vitro experiments do not necessarily translate into in vivo results. The small cross-interaction energies in FUS-PEG systems indicate condensate formation is driven by weak, non-specific attractions. For the G3BP1-PolyA system, we found that on average 4 PolyA molecules are present for every G3BP1 molecule. However, because PolyA has a much lower molecular weight than G3BP1, the majority of polymers in the condensate by mass and volume is instead G3BP1. The G3BP1-PolyA system has strong, specific attractions between polymers, rendering Flory-Huggins theory inapplicable in this scenario and more detailed models are needed to qualitatively explain the results.

We further note that to give a quick assessment of the sign of the tie-line gradient only two point measurements are needed in principle. One can first prepare a phase-separated sample and measure the dilute phase concentration of one solute, and prepare another with higher total concentration of that same solute while keeping other conditions constant. If the dilute phase concentration increases after adding the solute this indicates a positive tie-line and vice versa, while a constant dilute phase concentration simply corresponds to a flat tie-line. Similar linescans can be performed in this direction as well.

## IV. CONCLUSION AND OUTLOOK

We have provided a theoretical framework for connecting tie-lines to molecular interactions in the study of LLPS, particularly in multicomponent biopolymer systems which are of high interest in many areas of biology. Measurement of tie-line gradients allows for a quantitative description of stoichiometry and nature of interactions between solutes in a ternary system. Since biological condensates contain multiple species of proteins and nucleic acids, there is a clear need for quantitative biophysical characterisation of the roles that different components play in the LLPS process and our approach provides a method for parsing the interactions apart.

As an application of the theoretical framework, we developed an experimental method using microfluidics and a home-built photon-counting confocal which allowed us to monitor the dilute phase concentration of a phase separating protein, and therefore determine tie-lines by carrying out linescans across concentration series. Our findings highlighted that the protein FUS forms associative interactions with PEG with a high stoichiometry of PEG present in the condensates, and therefore contradicts previous claims that PEG is an inert crowder. Additionally, we studied the biomolecular system of G3BP1-PolyA RNA and found that our measurements agree with previous results of associative interactions between G3BP1 and RNA. We further discovered that the condensates tested had a G3BP1:PolyA RNA molar ratio of 1:4. Quantitative fitting of these gradients is however challenging due to the mean-field nature of the Flory-Huggins model as well as difficulties in solving for the exact binodal. It is worth mentioning that more detailed models exist that account for strong binding between solutes such as the sticker-and-spacer model [28] and phase diagrams with tie-lines can be computed via simulation [29]. Quantitative fitting of tie-line gradients is however beyond the scope of this paper. Having developed this robust theoretical and experimental framework for quantifying biopolymeric interactions in LLPS systems, many more systems may be studied to gain further mechanistic insight to the driving forces behind LLPS. Once interactions between different components are further understood, one may be able to better design therapeutics to selectively disrupt or enhance LLPS of disease-relevant biomolecules with greater mechanistic insight.

## Author Contributions

D.Q, T.J.W, N.A.E., T.C.T.M. and T.P.J.K. conceived the study. D.Q, T.J.W. and N.A.E. performed investigation. S.A., J.N.A., G.K., T.P.J.K. and P.St.G.H. provided resources. T.P.J.K. and P.St.G-H. acquired funding. D.Q., T.J.W. and N.A.E. wrote the original draft, all authors reviewed and edited the paper.

## Appendix A

### Mathematical details

#### 1. Minimal Flory-Huggins model

The free energy density in the minimal model is *f* (*ϕ*_1_, *ϕ*_2_) = (1 − *ϕ*_1_ − *ϕ*_2_) ln(1 − *ϕ*_1_ − *ϕ*_2_) + *ϕ*_1_ ln *ϕ*_1_ + *ϕ*_2_ ln *ϕ*_2_ + *χϕ*_1_*ϕ*_2_, and *ϕ*_1_, *ϕ*_2_ have to fall in the physically allowed region 0 *< ϕ*_1_, 0 *< ϕ*_2_ and *ϕ*_1_ + *ϕ*_2_ *<* 1. We first calculate the critical values of *χ* -where the spinodal region is a single point - and then calculate the critical points where the spinodal and binodal meet.

The spinodal boundary satisfies the condition det*H*_*µν*_ = 0 while within the physically allowed region. To relax the constraint we instead treat the mixing energy as Gibbs free energy and use number of particles as natural variables. This gives instead

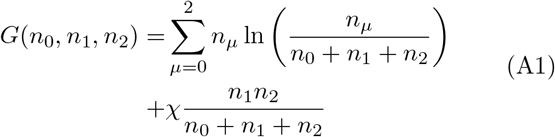

and the critical instability condition is that any one of the three chemical potentials have a vanishing second derivative. This condition, instead of a vanishing Hessian, arises because the three components are decoupled and the Hessian of *G* is always 0. The constraints are simply *n*_*µ*_ ≥ 0.

In the case 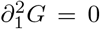 where 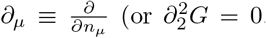, these give the same expression): (*n*_0_ + *n*_1_ + *n*_2_)^2^ − 2*χn*_1_*n*_2_ = 0. Solving for *n*_1_ and *n*_2_ in terms of *n*_0_, subject to the condition that only one set of solution exists we get *n*_0_ = *n*_0_, *n*_1_ = (*χ* − 1)*n*_2_ − *n*_0_ and 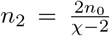. A critical interaction parameter 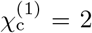 can be identified from the condition *n*_2_ *>* 0. This value corresponds to the critical interaction parameter for 2-component Flory-Huggins free energy. To calculate the volume fractions at 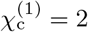 we first re-write *ϕ*_*µ*_ = *n*_*µ*_*/*(*n*_0_+ *n*_1_+ *n*_2_) and then substitute *χ* = 2. This gives

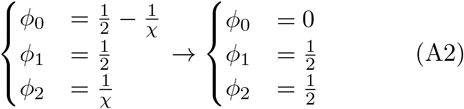

The concentrations thus fall onto the boundary of the physically allowed region and this case becomes intractable if we were to use the Hessian and apply the physical constraint.

For 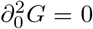 we get an equation quadratic in *n*_0_ with discriminant 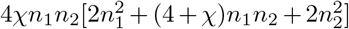. Setting it to zero, we obtain another quadratic equation in *n*_1_ with discriminant 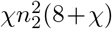 from which we obtain 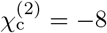. The concentrations at this point are:

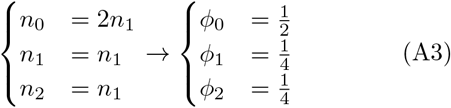

Since the second critical interaction 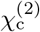 has concentrations all within the physically allowed region, we can extend this to general *N*_1_, *N*_2_ ≠ 1 using the Hessian. Starting with the equation det *H*_*µν*_ = 0 and setting the numerator to zero we obtain a polynomial quadratic in both *ϕ*_1_ and *ϕ*_2_. Taking the discriminant twice with respect to *ϕ*_1_, *ϕ*_2_ and setting the result to 0 gives

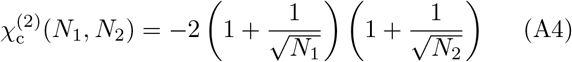

and the corresponding concentrations are

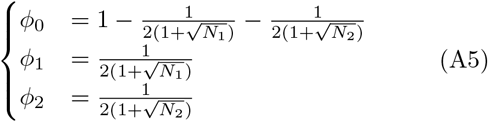

It is interesting to note the functional form of 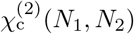 is very similar to that for the general 2-component system:

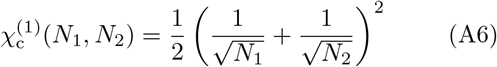

Next we calculate the critical points where spinodal and binodal meet. The interaction parameter *χ* has to satisfy *χ >* 2 or *χ <* −8 so the spinodal and binodal exist and are not single points. Since the spinodal is completely contained within the binodal region and both curves are smooth, they are co-tangent to one another at critical points. The tangent of the spinodal can be worked out easily by differentiating the equation 0 = det *H*_*µν*_ = *χ*^2^*ϕ*_1_*ϕ*_2_(1 − *ϕ*_1_ − *ϕ*_2_) + 2*χϕ*_1_*ϕ*_2_ − 1 and gives

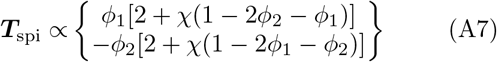

For the tangent of the binodal, we observe that at a critical point the ends of tie-lines shrink to a single point and thus must be asymptotically parallel to the direction of least stability, which is exactly the eigenvector that corresponds to the smaller of the eigenvalues of *H*_*µν*_. We thus obtain

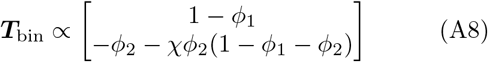

The critical points satisfy ***T***_spi_ × ***T***_bin_ = 0 as well as the spinodal equation (since it is on the spinodal), and the two equations have to be solved simultaneously to determine the critical point positions. Using *Mathematica* these can be solved to give the following points, subject to the constraint that they have to lie within the physically allowed regions:

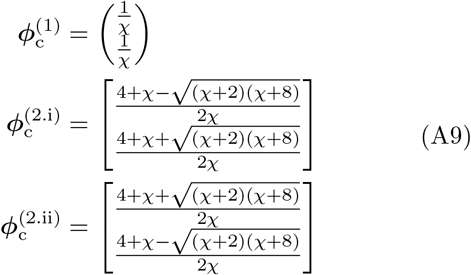

#### 2. Tie-line direction

The Hessian is defined as 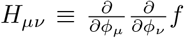 and using the Flory-Huggins expression it is

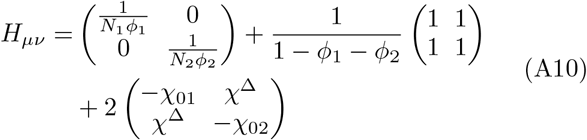

where 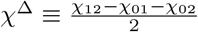. The first term is entropic contribution from solutes, which decouples the *ϕ*_1_ and *ϕ*_2_ directions due to its diagonal form; the second term is entropic contribution from the solvent and it maximally mixes the two directions; the third term has the form of an interaction matrix and its mixing/demixing tendency is a competition between the manifest self-energies −*χ*_01_ and −*χ*_02_ (in the diagonal position) and the cross-energy *χ*^Δ^ (in the off-diagonal position). In the dilute regime we approximate 1 − *ϕ*_1_ − *ϕ*_2_ ≈ 1 and absorb the solute entropy into the interaction parameters, redefining the interaction matrix by writing 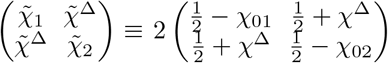. Furthermore we absorb the segment numbers *Nμ* into *ϕμ* as well: 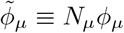. The Hessian becomes 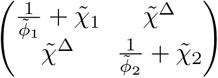. The eigensystem for the simplified *H*_*µν*_ can be easily worked out and we identify the eigenvector corresponding to the smaller eigenvalue as a proxy for the tie-line direction, since it is the direction of least free energy increase for phase separation. The gradient of this eigenvector is

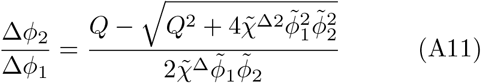

with 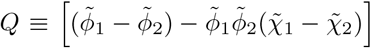. The numerator is always negative so the sign of this gradient completely depends on 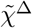, we thus claim 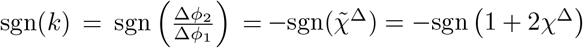. In experiments we observe that the amount of proteins used corresponds to a very small volume fraction, so we make the approximation 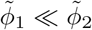 without loss of generality. Keeping only zeroth order terms in the numerator we arrive at the approximate gradient

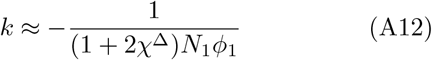

It is interesting to note that the homotypic interactions have totally disappeared.

On the other hand, if one only has information on the phase boundary in the dilute regime the sign of 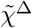 is inaccessible. To demonstrate the point we calculate the spinodal boundary by setting the Hessian at low *ϕ*_1_, *ϕ*_2_ to zero, obtaining

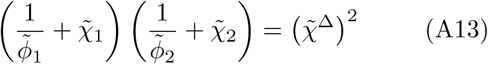

As such, the spinodals near the origin do not depend on the sign of 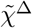 at all. Although phase diagrams measured in experiments are assumed to be the binodal, we assume that the binodal shape is not qualitatively different from the spinodal so the experimentally measured dilute phase boundary alone is not sufficient to decipher the mechanisms of phase separation.

## Appendix B

### Method and materials

#### 1. Home-built setup

##### Photon-counting setup

Measurement of tie-lines requires a robust method for measuring dilute phase concentrations of proteins in heterogeneous mixtures of condensates and soluble monomer. In order to carry out these measurements, we utilised a droplet microfluidic technique in which we created microdroplets containing the desired concentrations of protein and co-polymer and then measured these droplets on a home-built confocal setup [25]. In brief a 488 laser line is coupled into a 60x-magnification water-immersion objective (CFI Plan Apochromat WI 60x, NA 1.2, Nikon) and emitted photons are directed onto an avalanche photodiode (APD), a motorised XYZ stage is mounted on top of the objective so that a microfluidic device may be monitored throughout the experiment. A four-inlet microfluidic device is used to generate water-in-oil droplets of varying concentrations of solutes, with one inlet connected to an oil syringe and three inlets to syringes loaded with individual solutes or a dilution buffer. After being made, droplets were squeezed into a measurement channel and the actual measurement were made at the end of the channel to give sufficient incubation time (60-80s) for the system to equilibrate. The confocal setup is illustrated in figure 4A and the microfluidic setup in figure 4B. The signal from the APD is binned into 1ms intervals to give out a photon intensity value in photons per sec (MHz), which is directly correlated with the concentration of protein so long as the protein is in the 1-10µM range where solution quenching may be neglected. The intensity trace typically has a baseline intensity (corresponding to dilute concentration of protein) and spikes corresponding to condensates passing through the confocal spot. These spikes are filtered out using a 3-*σ* cutoff. We measured intensity profiles of ∼ 100 droplets at each concentration composition and the baseline intensity of each droplet was calculated. The intensities were binned to give an overall signal estimate (figure 4C). To map the intensity to actual concentrations we performed calibration series without LLPS, and obtained monotonic conversion curves (figure 4D). As far as tie-line gradients are concerned, however, this last step of concentration calibration is not necessary since two equal intensity signals directly imply equal concentrations due to the one-to-one nature of conversion curves.

**FIG. 4.**
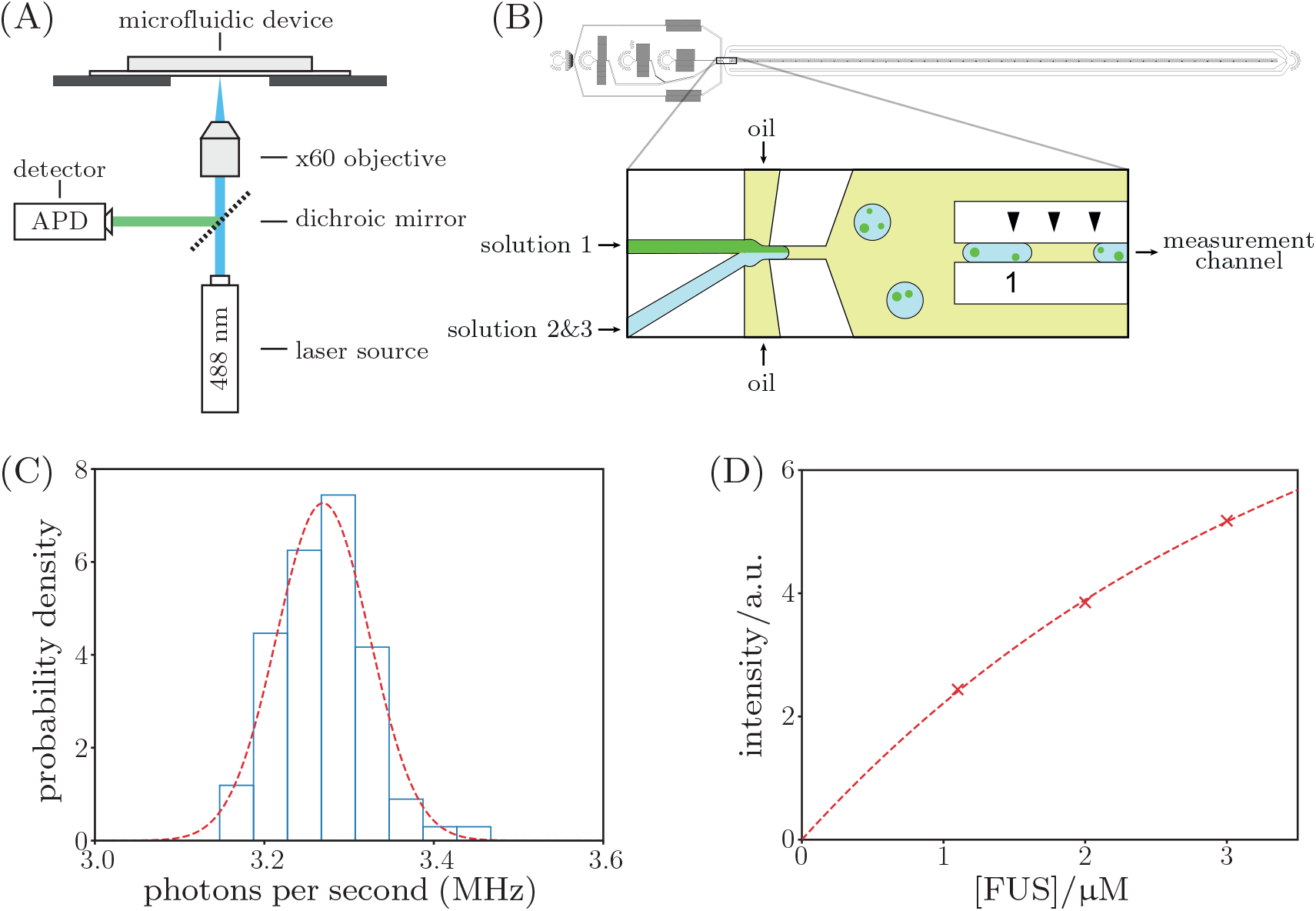
Experimental overview. (A) simplified illustration of the home-built confocal measurement setup, described in detail elsewhere [25]; (B) microfluidic device used, with a zoomed-in view of the droplet-generation section. (C) histogram of mean baseline signal for individual droplets, the red dashed line is a gaussian distribution using the mean and standard deviation calculated for the series of droplet. (D) calibration curve for correlating signal to actual concentration. The data (red crosses) was fitted to a quadratic curve (dotted line).

##### Microfluidic device fabrication

The microfluidic device was designed using AutoCAD software (Autodesk) and printed on acetate transparencies (Micro Lithography Services). The replica mold for fabricating the device was prepared through a single, standard soft-lithography step [30] by spinning SU-8 3025 photoresist (MicroChem) onto a polished silicon wafer to a height of around 25 µm. The UV exposure step was performed with a custom-built LED-based apparatus [31]. The mold was then used to generate a patterned PDMS slab. To this effect, the mold was casted in a 10:1 (w/w) mixture of PDMS (Dow Corning) and curing agent (Sylgard 184, Dow Corning), degassed and baked for 1.5h at 65°C. The formed PDMS slab was cut and peeled off the master and access holes for the inlet tubes were introduced using biopsy punches. The devices were then bonded to a thin glass coverslip after both the PDMS and the glass surface had been activated through oxygen plasma (Diener electronic, 40% power for 30s). After bonding, 1.5% trichloro(1H,1H,2H,2H-perfluorooctyl) silane (Sigma-Aldrich) in HFE7500 (Fluorochem) was injected into the PDMS devices to render their surface hydrophobic. After 5 minutes incubating in the silane solution, the solution was washed out with HFE7500 and the devices were heated at 95°C to remove any excess oil.

#### 2. Experimental conditions

##### FUS-PEG

FUS proteins used in our experiments are fluorescently tagged by eGFP. The FUS-eGFP protein is stored at 100uM in 50mM TRIS-HCl, 1M KCl under -80°C. To run the microfluidic experiments 3 separate syringes containing FUS, PEG, and buffer were prepared. Table II lists the content of each solution. A fourth syringe containing HFE-7500 oil (FluoroChem) and 1.2% PTFE-PEG surfactant (RAN Bio) was additionally prepared. Initial protein solution concentrations are measured with NanoPhotometer (IMPLEN, GENE-FLOW NP80) at 488nm wavelength.

**TABLE II.**
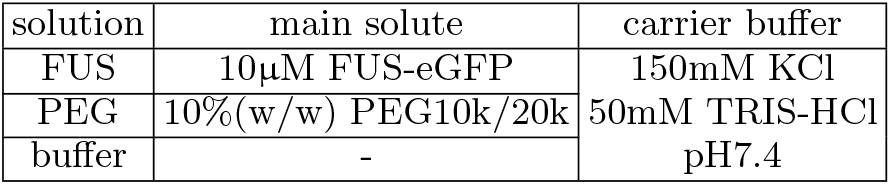
Solution content of syringes used in microfluidic experiments for FUS-PEG systems.

##### G3BP1-PolyA RNA

G3BP1 proteins used in our experiments are florescently tagged by mEmerald. Table III lists the content of solutions used in the G3BP1-PolyA RNA experiments. The fourth syringe contains the same HFE-7500 oil (FluoroChem) and 1.2% PTFE-PEG surfactant (RAN Bio) as before. Initial protein solution concentrations are measured with NanoPhotometer (IMPLEN, GENEFLOW NP80) at 488nm wavelength.

**TABLE III.**
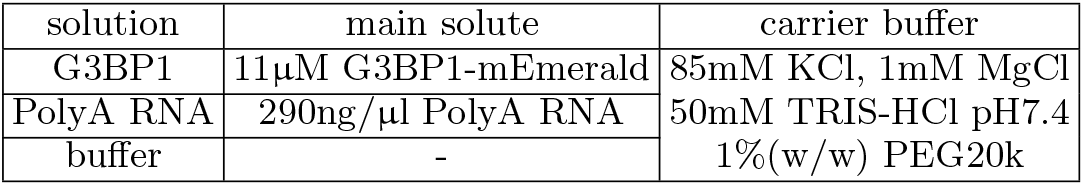
Solution content of syringes used in microfluidic experiments for the G3BP1-PolyA RNA system.

#### 3. Protein purification protocol

##### G3BP1 protein

Baculovirus expression system with Sf9 insect cells was used to express the recombinant pACEBac2 vector containing the cleavable (His and MBP tag) His-Emerald-G3BP1-MBP construct. Three days post viral infection the cell culture was harvested and the pellet was resuspended in Buffer A (50mM Tris, 1.0 M KCl, 1 mM EDTA, pH 7.5) containing protease inhibitors and 0.1% CHAPS. Using Dounce homogeniser, cells were lysed and the cell lysate was further clarified by centrifugation at 100,000g. Supernatant containing the overexpressed protein was subjected to a three step chromatography purification scheme. Ni-advance column was used to capture the bulk of the fusion-G3BP1 protein. The G3BP1 containing fractions, assessed by SDS-PAGE, were pooled and applied to an amylose column (NEB) for further purification before being treated with the 3C Precission Protease (Thermo Fisher) for the cleavage of the His and the MBP tags. Cleaved tags were further removed by applying the sample on a Superdex-200 Increase (Cytiva) size exclusion chromatography column in the storage buffer (50 mM Tris, 300 mM KCl, 1 mM DTT, pH 7.5 buffer). Pure fraction of Emerald-G3BP1 were pooled, concentrated and snap frozen in liquid nitrogen and stored at -80°C for the droplet assays.

##### FUS protein

Fluorescently tagged FUS protein used in this study was expressed and purified from an insect cell expression system following the previously published protocol [32].The purified protein was stored in the final storage buffer, 50 mM Tris, 1 M KCl, 1 mM DTT, 5% glycerol, pH 7.4 at 100 µM protein concentration.

#### 4. Physical parameters used in calculations

The densities, molecular weights and segment numbers used in calculations for various components are listed in table IV. To calculate *N* we note that *N* represents the number of lattice sites a single molecule occupies, and we approximate the lattice volume as the volume of a single amino acid residue (density 1.35g/ml and MW 110Da).

**TABLE IV.** Physical parameters used in calculations.

| solute | density(g/ml) | MW(kDa) | $N$ |
| --- | --- | --- | --- |
| FUS | 1.35 | 107 | 909 |
| G3BP1 | 1.35 | 87 | 791 |
| PEG20k | 1.125 | 20 | 218 |
| PEG10k | 1.125 | 10 | 109 |
| PolyA | 1.6 | 3 | 23 |

#### 5. Dilute phase data processing

##### Anchored linescan

The phenomenological curve we used to fit the linescan data is

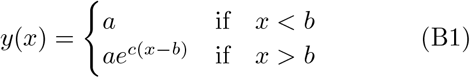

with *y* denoting the signal intensity and *x* the total concentration of the agent (in our case, PEG20k). *y*(*x*) has a plateau at low values of *x* and decays exponentially when *x* crosses some threshold *b*. Once we have this fit we can compare it to the anchor values *x*_0_, *y*_0_ (which we know) and work out *x*_1_ (which corresponds to the PEG20k concentration that gives the same intensity reading and thus same dilute phase FUS concentration) by writing *y*_0_ = *y*(*x*_1_). We then have 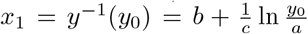 and Δ[PEG20k] = *x*_0_ − *x*_1_.

##### Serial linescan

To extract Δ[PEG10k] we again fit the data to equation (B1). For the two linescans we fix *a* to the total [FUS], treat *c* as a global fitting parameter and *b*’s fitting parameters for individual linescans. This allows us to extract Δ[PEG10k] in the following manner. Denote the fitted curves as *y*_*i*_(*x*) with parameters *a*_*i*_, *b*_*i*_ and *c*, with *i* = 1, 2. The condition that *y*_1_(*x*_1_) = *y*_2_(*x*_2_) simply gives ln *a*_1_ + *c*(*x*_1_ − *b*_1_) = ln *a*_2_ + *c*(*x*_2_ − *b*_2_) so 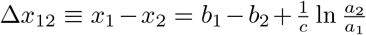, and it is essential that *c* is the same for both curves to allow the extraction to be performed. This assumes a constant tie-line gradient.

#### 6. Confocal microscopy

To check the direction of the tie-line, we imaged condensates with fluorescently labelled biopolymers using a Stellaris 5 confocal microscope equipped with 63x oil immersion objective (Leica HC PL APO 63X/1.40 OIL CS2, NA 1.4). The sample solution contained 3µM FUS and 6%(w/w) PEG20k, of which 0.5%(w/w) is fluorescently labelled. The samples were imaged in airtight imaging wells made from polydimethylsiloxane (PDMS) (Sylgard 184 kit; Dow Corning) in between 24×60 mm No.1.5 cover glass slides on the bottom (DWK Life Sciences) and 18×18 mm glass slides on top (Academy). Both glass and PDMS was treated with PEG(5000)-Silane (Sigma-Aldrich) by submerging them in a solution of 5 mg of PEG(5000)-Silane in 20 µl of 50% acetic acid and 1 mL of ethanol for 1 hour at 65°C. Wells and glass were washed rigorously before use with water in a sonication bath.

## Appendix C

### Numerical fitting

Using the FUS-PEG10k data we re-constructed the phase diagram and tie-lines, and used the Flory-Huggins free energy to fit the interaction parameters. The results are shown in figure 5. The fitted interaction parameters] are *χ*_FUS-water_ = 0.530, *χ*_PEG-water_ = 0.602 and *χ*_FUS-PEG_ = −0.063, giving 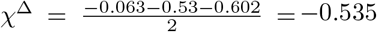. This is not far from the *χ*^Δ^ = −0.74 calculated using the Hessian.

**FIG. 5.**
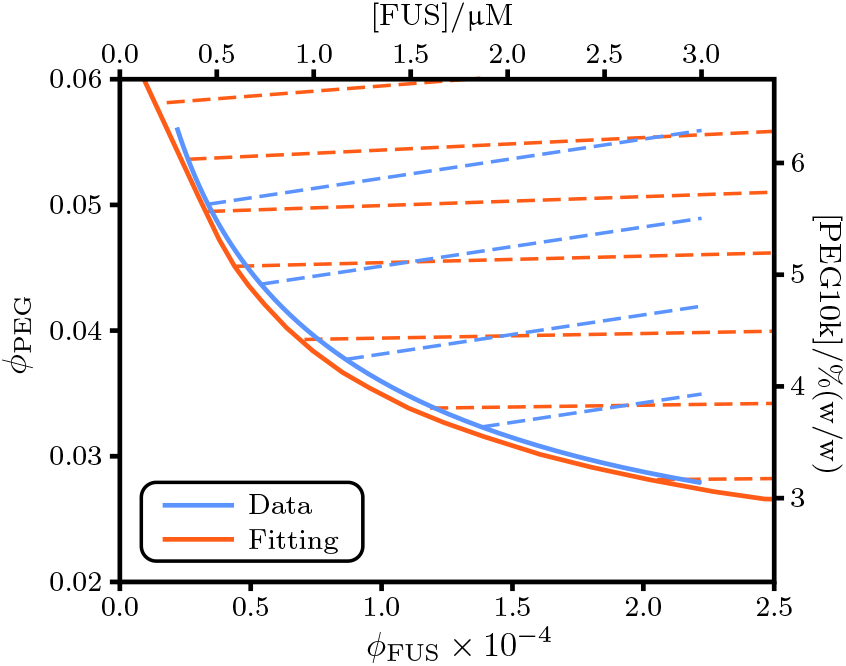
Full linescan data for FUS-Dextran system.

## Appendix D

### FUS-Dextran results

In addition to PEG, Dextran (500kDa) is often used to induce phase separation of FUS [33]. To test whether Dextran was an inert crowder or indeed an associative polymer as we carried out linescan experiments on FUS vs dextran. Sample conditions are the same as that used in FUS-PEG systems. Our findings show that as FUS concentration is increased, the dilute concentrations are positively shifted at the same dextran concentration and that the dilute concentrations are similarly decreasing as dextran is increased for a single FUS concentration (figure 6). This is the same behaviour observed for PEG10K and 20K, and thus concludes that dextran behaves as an associative polymer during its role in FUS phase separation.

**FIG. 6.**
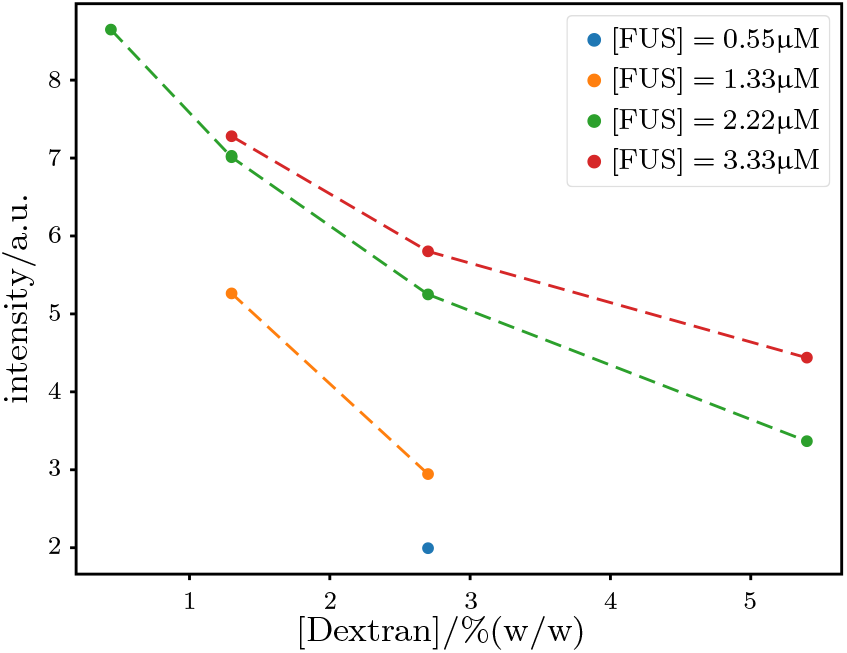
Full linescan data for FUS-Dextran system.

